# Effect of *Geobacter metallireducens* nanowire on electron transfer efficiency in microbial fuel cell

**DOI:** 10.1101/2021.07.14.452433

**Authors:** Shunliang Liu, Yali Feng, Haoran Li

## Abstract

The inhibitory effect of electron mediator 2,6-anthraquinone disulphonate (AQDS) on *Geobacter metallireducens* nanowire in the microbial fuel cell (MFC) was studied. In the culture process of *G.metallireducens* with Fe(OH)_3_ as an electron acceptor, the concentration of reduction product Fe (II) in solution without AQDS was higher than that with AQDS after 10 days, due to the formation of microbial nanowires. The effects of nanowire on electron transfer efficiency and electrical current characteristic were studied using a double chamber MFC reactor. The transfer efficiency between biofilm and electrodes was increased by nanowire, which increased the maximum output voltage of MFC was 442 mV. The nanowire biofilm electrode had a bigger cyclic voltammetry curve peak, smaller activation resistance, and a stronger current response signal through electrochemical measurement, which indicates that the nanowire enhanced the electrochemical activity of the electrode.

## 1 Introduction

Microbial fuel cells (MFC) can achieve both purposes of wastewater treatment and electrical energy production, through converting the chemical energy of organic compounds into electrical energy using microorganisms as catalysts (Hu et al., 2017). Because of the advantages of extensive substrates sources, easy reaction conditions, efficient processing capability, MFC have been widely researched to treat waste-water such as organic (Asefi et al., 2019; Priya et al, 2019), cyanide (Wu et al, 2019), and high-price metal (Li et al. 2019). At present one of the limitations of MFC technology is the low output power, and various means have been used to improve the efficiency of electron transfer (S. Kalathil et al, 2018). For instance, the development of MFC electrode materials includes carbon felt (Feng et al, 2019), graphite foam (Chen et al, 2019), metal material (Liu et al, 2018) and natural biomass electrode material; Modification of electrode materials, such as graphene-modified electrode (Lin et al, 2019), non-metallic (Bajracharya et al, 2019) and metal element doped electrode (Palanisamy et al, 2019), carbon nanotubes modified electrode (Delord et al, 2019), conductive polymer (polyaniline (Zhai et al, 2019), polypyrrole (Anders et al, 2009), silane coupling agent) modified electrode materials; Adding electronic mediators such as humic acid (Anders et al, 2009), anthraquinone compounds (Tang et al, 2017), mercaptan-containing molecules, cysteine, melanin (Costa et al, 2009) to improve the electron transfer efficiency. Among them, quinone compounds are the preferred electronic medium in the process of biodegradation, which can not only transfer electrons in the microbial reaction and act as the REDOX medium but also the hydroquinone decomposition products can further participate in the subsequent pure chemical reaction. Many studies have confirmed that AQDS acts as an electron shuttle, favoring the transfer of electrons between microorganisms and organic matter (Lovley D R et al, 1996; Luu et al, 2003), and also between electron donors and electron acceptors (Yang et al, 2009). Evidence from other studies suggests that only a small amount of AQDS (0.01-0.20 mM AQDS) is enough to “shuttle” electrons between microorganisms and oxides, nanocilia-like PANI grew uniformly on the surface of rGONF under the guidance of the low-level AQDS to improve successfully the specific capacitance and electrochemical stability of asymmetric supercapacitor (ASC) (Du et al, 2021), moreover positive effect that in the reductive dissolution of As(V)/Fe(III) during sediment supplementation with lower (0.05 mM) compared to the high level (0.10 mM) of AQDS, whereas an inhibitory effect resulting from even higher (1.00 mM) level of AQDS (Chen et al, 2017). However, at present, AQDS as an electronic medium to improve the efficiency of MFC electron transfer is less influential (Santoro et al, 2017). Nevertheless, how the electron shuttle compounds affect MFC electron transfer behavior and processes of the fabricate nanowire by microorganisms remains unknown.

The theory of nanowire transfer in MFC refers to that some microorganisms produce a conductive pilin protein (nanowires) in the process of growth and metabolism, and the electrons can be transferred to the anode electrodes through nanowires (N. Alves et al, 2016). The microbes are connected to the electrodes through nanowires, thus releasing the transmission constraints between cell membranes and electrodes. Not only the biofilm on the surface of the electrode can transfer electrons to the electrodes but also the outer microbes can transfer electrons through nanowires, which can increase the output current of MFC (Numfon et al, 2016). At present, many bacteria have been observed such as Geobacterium, Schwarzschild, Shewanella oneidensis, Synechocysti, *Pelotomaculum thermopropionicum* producing nanowires (Ganesh et al, 2017). Among them, the Geolimetal Metallireducens bacteria are very efficient in remote extracellular electron transfer, which can effectively explain the bridging relationship between cytoplasmic electron donor and extracellular receptor, and better demonstrate the influence of nanowires electronic media (Marisa et al, 2020).

To improve the electronic transfer efficiency of microbes and the output voltage of MFC, the generation mechanism of *Geobacter metallireducens* nanowire by the electronic mediator (AQDS) has been studied. Meanwhile, the effect of nanowires on the electron transfer speed, the metal reduction efficiency, and output voltage was explored with a double chamber MFC device. The effect of nanowire on MFC efficiency by electrochemical means was examined. This study investigated the inducing effect of insoluble electron acceptors on nanowires generation, which will be very important for understanding MFC, soil remediation, and solid-phase fermentation using solid-phase as electron acceptor.

## 2 Materials and methods

### 2.1 Growth medium and microorganism

*Geobacter metallireducens* (*G.metallireducens*), a German Collection of Microorganisms and Cell Cultures preserved strain (DSMZ 7210, ATCC 53774), was used as an experimental strain. The composition of the synthetic medium used in the study was: sodium acetate (10 mmol/L), KCl (0.1 g/L), NH_4_Cl (0.2 g/L), NaH_2_PO4 (0.6 g/L), NaHCO_3_ (2.5 g/L), Wolfes’ trace mineral solution (10 mL/L) and Wolfes’ vitamin solution (10 mL/L), pH 6. 8~7. 0. The gas N_2_-CO_2_ (8:2) was used to remove oxygen from the growth medium, then the growth medium was separated into anaerobic culture tubes to be sterilized at 121 °C for 15 min.

### 2.2 MFC construction and operation

The structure of the double chamber MFC reactor is shown in Fig.1. The reactor consists of 250 mL cathode and 250 mL anode chamber connected by a proton exchange membrane (Nafion-117, DuPont). The electrode was a pure graphite electrode with a surface area of 75 cm^2^, which was cleaned by 1.0 mol/L HCl and 1.0 mol/L NaOH to remove impurity ions and adsorbed microbes respectively. The mixed liquid with a ratio of bacterial solution to anolyte is 1:9 was added to the MFC anode chamber, and the mixture gas N_2_-CO_2_ (8:2) was slowly passed through to remove oxygen. The catholyte was added to the cathode chamber, and the air was continuously insufflated to maintain dissolved oxygen concentration. The anolyte contained KCl 0.1 g/L, NH_4_Cl 0.2 g/L, NaH_2_PO_4_ 0.6 g/L, NaHCO_3_ 2.5 g/L, NaCl 2.9 g/L, Wolfes’ vitamin solution 10 mL/L, Wolfes’ mineral solution 10 mL/L, electron donor NaAC 10 mmol/L. And the composition of catholyte is KCl 0.1 g/L, NH_4_Cl 0.2 g/L, NaH_2_PO_4_ 0.6 g/L, NaCl 2.9 g/L, Tric-HCl (adjust pH to 7.0). 50ummol/L and 0ummol/L of AQDS will be adding to the corresponding MFC and set 3 parallels for each concentration gradient under the experiment, respectively. The cell voltages were measured every 5 min using a data acquisition system (RBH8223H, Ribohua Co.) across an external resistance of 510 Ω. At the end of the MFC operation cycle, 10 mmol/L electron donor NaAc was added into the anode for the next cycle.

**Fig 1.**
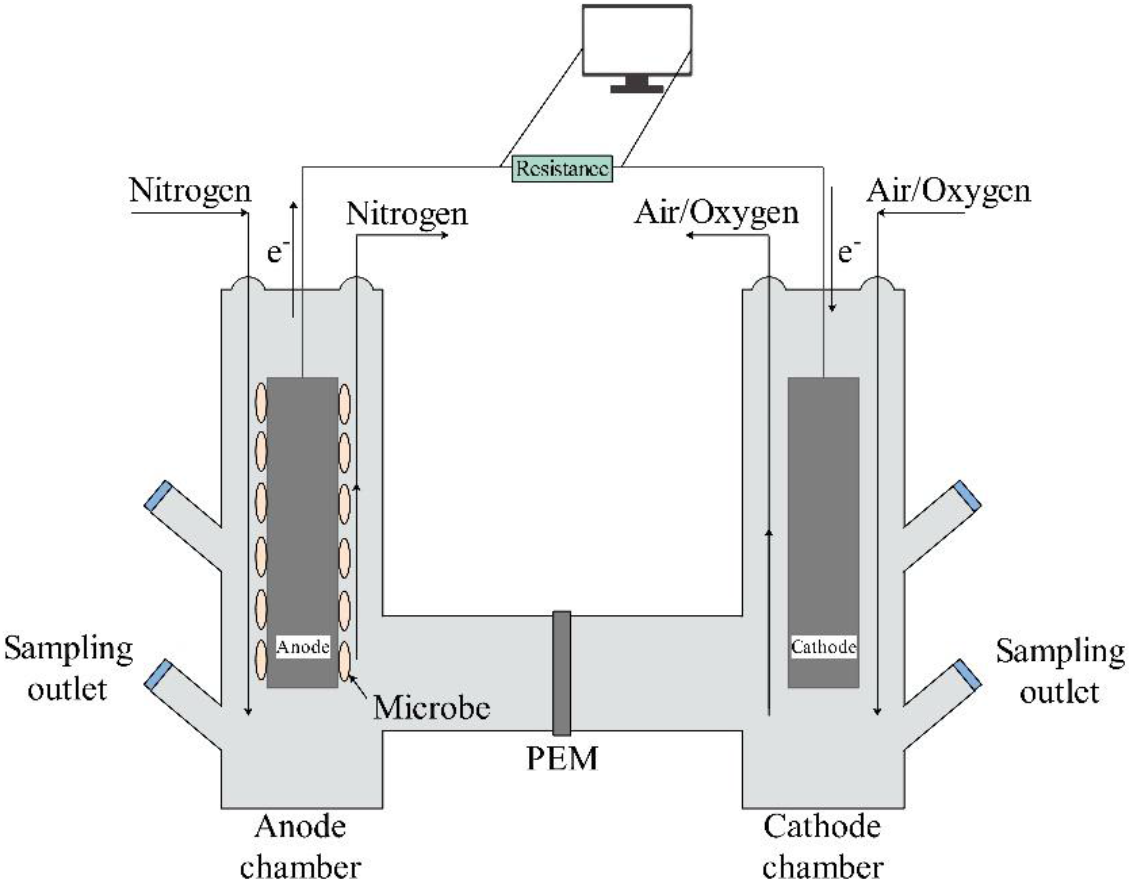
The structure of the double chamber MFC reactor

### 2.3 Electrochemical tests

Electrochemical measurements include cyclic voltammetry (CV), electro-chemical impedance spectroscopy (EIS) and linear sweep chronoamperometry (LSC) were carried by CHI660D electrochemistry tester with a three-electrode electro-chemical system. The electrochemical tests were researched using the MFC anode as the working electrode, a platinum electrode as a counter electrode, and a saturated calomel electrode as the reference electrode. In a mixed buffer (10 mmol/L phosphates +10 mmol/L NaAc), CV spectra were recorded at the scanning rate of 5 mV/s. And LSC spectra were recorded at the scanning voltage range of 0.8~0 V, scanning rate of 0.01 mV/s. In a 10 mmol/L K_3_Fe(CN)_6_/K_4_Fe(CN)_6_ (1:1) + 0.1 mol/L KCl solution, EIS was measured for MFC anode in a frequency range of 0.1Hz~100kHz with an AC signal of 5 mV amplitude.

### 2.4 Scanning electron microscopy

The microbial structure of enriched bacteria and biofilm on the MFC anode surface were examined using scanning electron microscopy (JSM-7001F, JEOL Ltd). The electrodes were fixed with 2.5% glutaraldehyde at 4 °C for 2~4 h, and rinsed 3 times with 100 mmol/L sodium cacodylate buffer (pH 6.8) for 10 min. Then fixed electrode was dehydrated gradually with 50%, 70%, 80%, and 90% ethanol for 15 ~ 20 minutes, and gently washed twice with pure isoamyl acetate for 15 min. The samples were placed in the critical evaporator to be dried for 4 h, and the dried samples were coated with gold prior to SEM analysis.

## 3 Results and discussion

### 3.1 Generation of bacterial nanowires

Metal reducing bacteria take metal oxides as the final electron acceptor to form a complete electron transfer chain through extracellular electron transfer, which is the metal reduction process by microbes. Ineffective extracellular electron transport will impact intracellular electron transport negatively, which prevents bacterium to synthesize adenosine triphosphate (ATP), the energy source of activities for metabolism and growth (Okamoto et al, 2014). The low bioavailability of electron acceptors leads to inefficient extracellular electron transport and inactive vital movement of bacteria. And the efficiency of bioleaching and environmental remediation is reduced (Liu et al, 2014).

Electron mediators act as electronic transport carriers in dissimilatory reduction system, which can accelerate the electron transfer rate between microbes and solid oxides. For example, intracellular reduction cytochromes lose electrons to free AQDS (9,10-anthraquinone-2,6-sulfonate) to form AHQDS molecules. The AHQDS containing electrons moved to the surface of Fe(OH)_3_ for transmitting electrons. Then by losing electrons AHQDS convert back to AQDS to transport electrons for the next round (Pruetsaji et al, 2018; Han et al, 2017). The process is shown in Fig. 2a. The equation is as following:

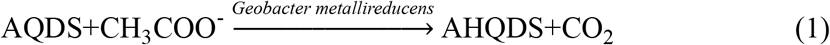

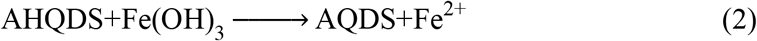

**Fig.2.**
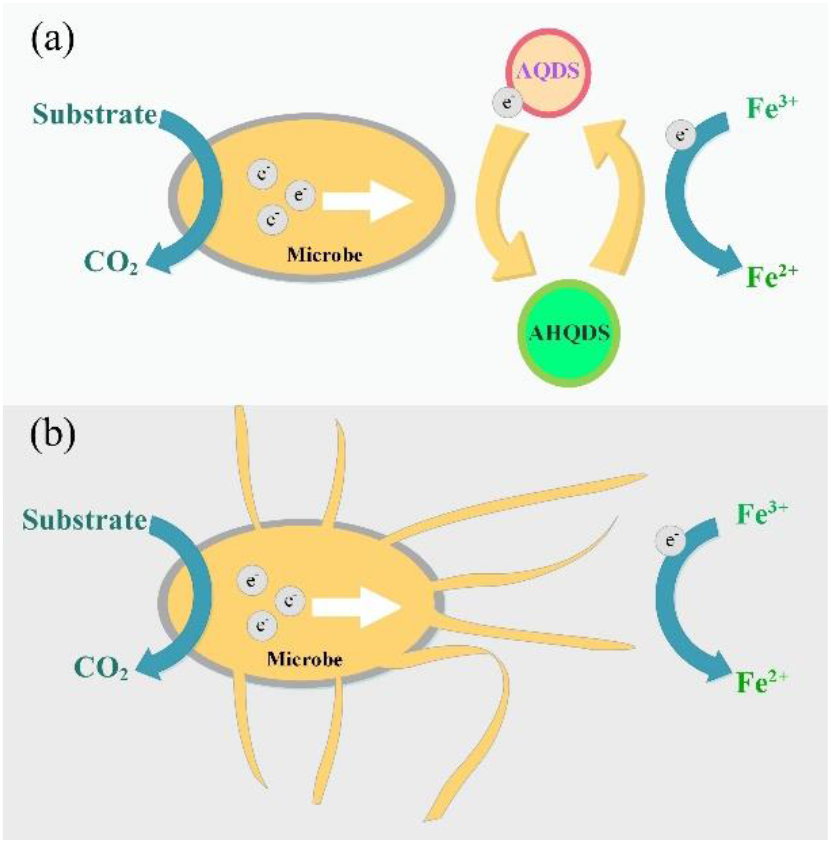
Electron transfer pathways for *G.metallireducens* during the reduction process

In recent years, it has been found that *G.metallireducens* can generate conductive nanowires similar to additional pili, due to the lack of soluble electron acceptors (Sébastien et al, 2017). Microbes transfer electrons far away from the cell surface through highly efficient conductive nanowires, thereby, microbes break through the electron transfer restriction which requires direct contact with solid electron acceptors or adds electronic mediators (Mónica et al, 2016). Microbial nanowires are an effective way that microorganisms evolve to improve the efficiency of extracellular electron transfer (Toshiyuki et al, 2018). The process is shown in Fig. 2b.

The growth of nanowires of reducing bacteria was inhibited by adding AQDS, and the effect of nanowires on the reduction efficiency was compared. Fig.3 shows nanowire production on the reduction efficiency of Fe(OH)_3_ (100mM) with an initial concentration of electron donor (NaAc) 10 mM.

**Fig.3.**
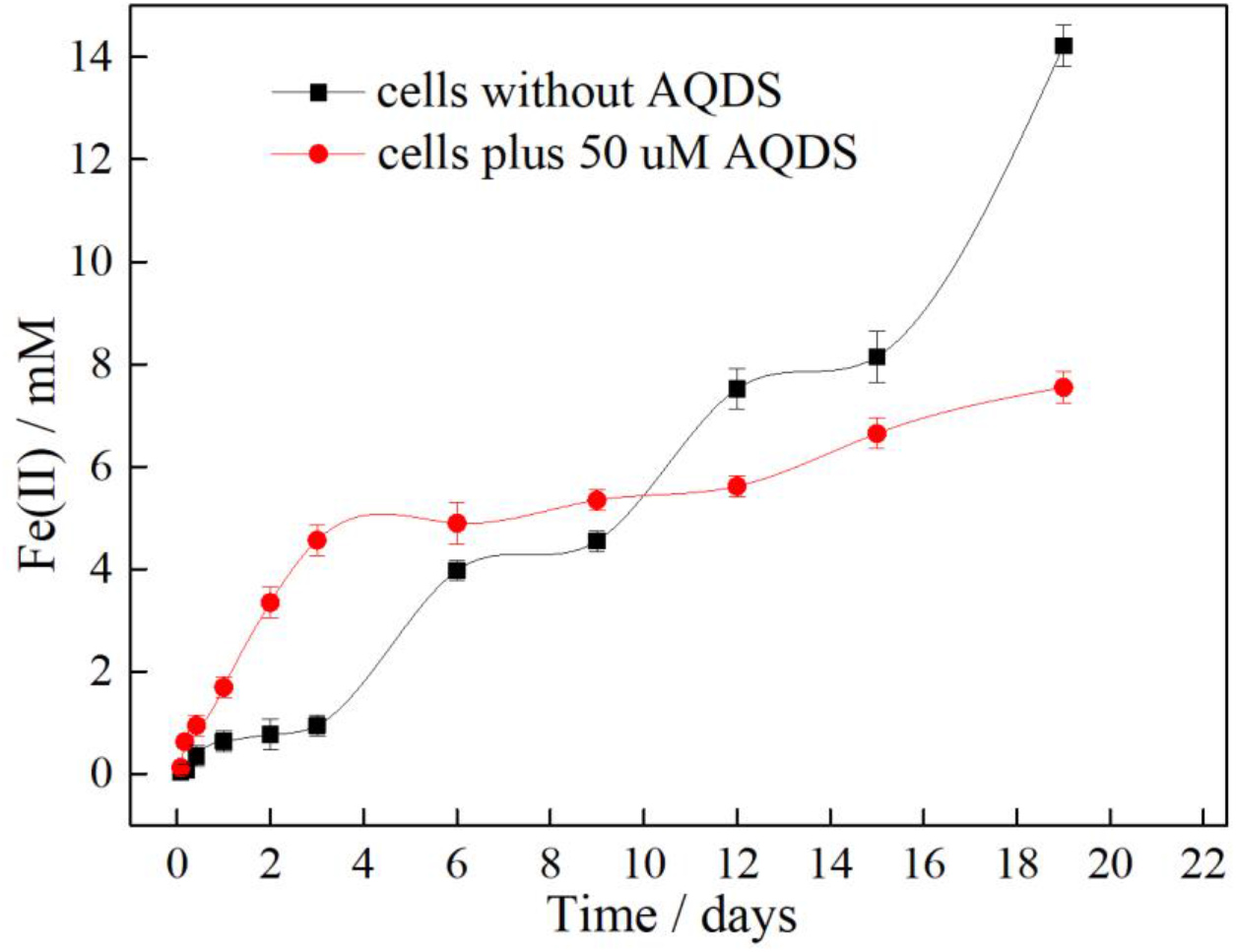
Effect of AQDS on the dissimilatory reduction of Fe(OH)_3_ (100mM)

It can be seen from Fig.3 that the reduction of Fe(OH)_3_ can be accelerated by AQDS in a short time (within 10 days) after inoculating microbes in the culture system. Compared with using Fe(OH)_3_ as a microbial electron acceptor directly, the addition of AQDS can avoid the extracellular electron transfer process. The AQDS would act as an electron carrier to shuttle between the cell membrane and metal hydroxide, thus accelerating the dissimilatory reduction of Fe(OH)_3_. However, the accelerating effect of AQDS on microbial reduction was not significant after a long reaction time (10 days). Microbes grow nanowires without AQDS for a long time, which promoting the electron transfer exceeds AQDS in the reduction process (the concentration of reduction product Fe(II) is higher than that with AQDS). The addition of AQDS hinders the growth of microbial nanowires. Although the electron transfer is accelerated in the initial stage, the upper bound of transfer efficiency is worse than that of nanowires. Fig. 4 shows the microbial structure of *G.metallireducens* in the solution (reaction for 20 days). It can be observed that the addition of electronic mediator AQDS inhibit the production of nanowires (Fig.4a), while in a solution without electronic mediator AQDS, *G.metallireducens* is stimulated to produce nanowires (Fig.4b).

**Fig.4.**
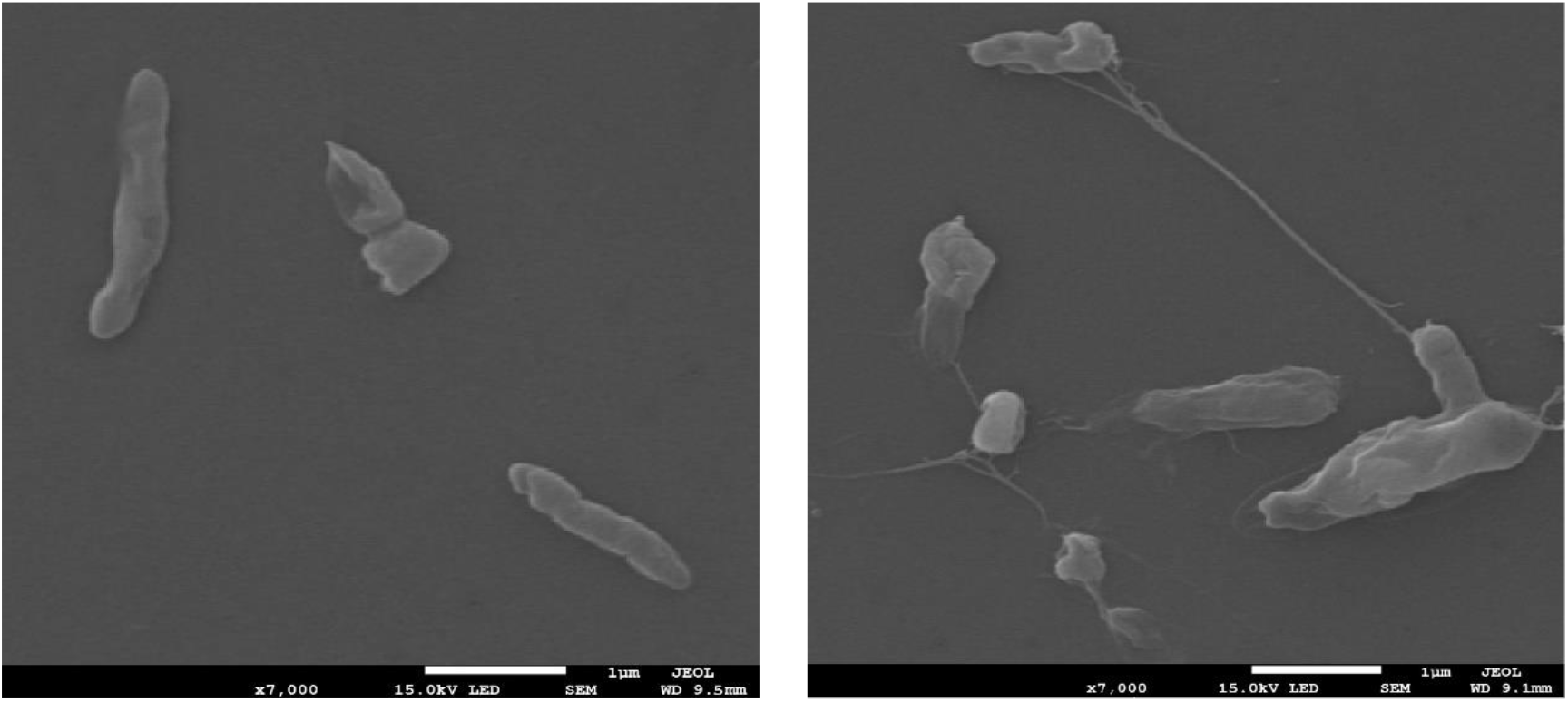
The microbial structure of *G.metallireducens* in the solution (reaction for 20 days)

### 3.2 Effect of nanowires in the MFC

To accurately research the effect of *G.metallireducens* nanowire on electron transfer efficiency, MFC is used to study the electron transfer process. Anodic graphite electrode was used as the final solid electron acceptor instead of Fe(OH)_3_. Electrons reduce dissolved oxygen by anodic graphite electrode-external circuit-cathode, which maintaining the electron transfer process from microbes to solid electron receptors. By recording the current value of the external circuit, the process of electron transfer can be accurately reflected. The reaction process of MFC is as following:

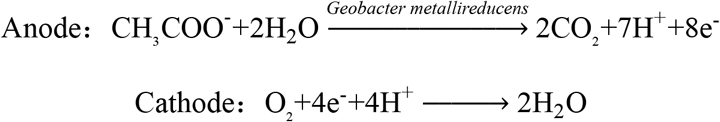

The electricity voltage output obtained from the MFC inoculating *G.metallireducens* (10%) is shown in Fig.5(a). As can be seen from Fig.5(a), MFC can be activated quickly with 50μm AQDS, and the maximum voltage of MFC can be stabilized at about 500 mV. MFC was activated slow Without AQDS, and the maximum voltage of MFC was stabilized at 398 mV. Adding electronic mediator AQDS in starting period of MFC can significantly increase the output voltage of MFC, and AQDS acts as a transfer electron role between cells and solid electron acceptors. However, electron transfer efficiency of MFC without AQDS is low, due to the incomplete nanowires.

**Fig.5.**
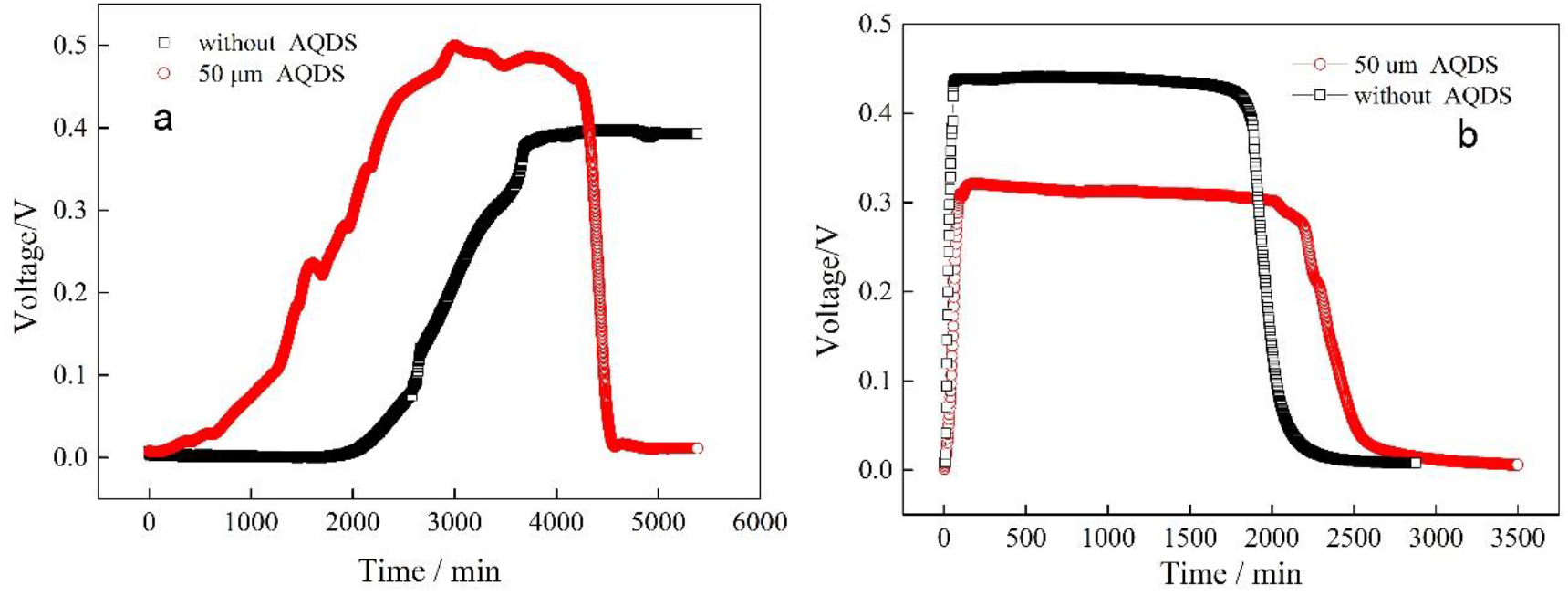
Electron current in a microbial fuel cell by *G.metallireducens*. (a)In the microbial fuel cell, graphite electrode served as the sole electron acceptor, acetate as electron donor, 10% inoculum. (b) Voltage output of MFC which solution replaced after operation stability (10d)

To exclude the effect of AQDS on MFC, and study the relationship between microbial nanowire with electron transfer, the solid electron acceptor (anode graphite electrode) and the culture solution were separated. The new acetate solution was used to replace the culture solution in the anode chamber, the graphite electrode which formed the biofilm was retained, and then the MFC was reconnected. The Voltage output of the MFC solution was replaced after operation and stability were indicated in Fig.5(b). The output voltage of MFC without AQDS has a slight change, and the maximum output voltage was 442 mV. However, due to the removal of AQDS, the maximum output voltage of MFC with AQDS decreased significantly from 498 mV to 321 mV. The results showed that the electron transfer efficiency of nanowire biofilm is higher than that of biofilm without nanowires.

Fig.6 shows MFC anode biofilm with AQDS (Fig.6a) and without AQDS (Fig. 6b). It can be observed by comparing the figure that the *G.metallireducens* did not generate nanowires in biofilm with AQDS. Microbes adhere to the graphite fibers of electrodes such as granules. And the *G.metallireducens* generated nanowires in biofilm without AQDS. Nanowires are interlaced to connect microbes in biofilm, or microbes adhere to graphite fibers through nanowires.

**Fig.6.**
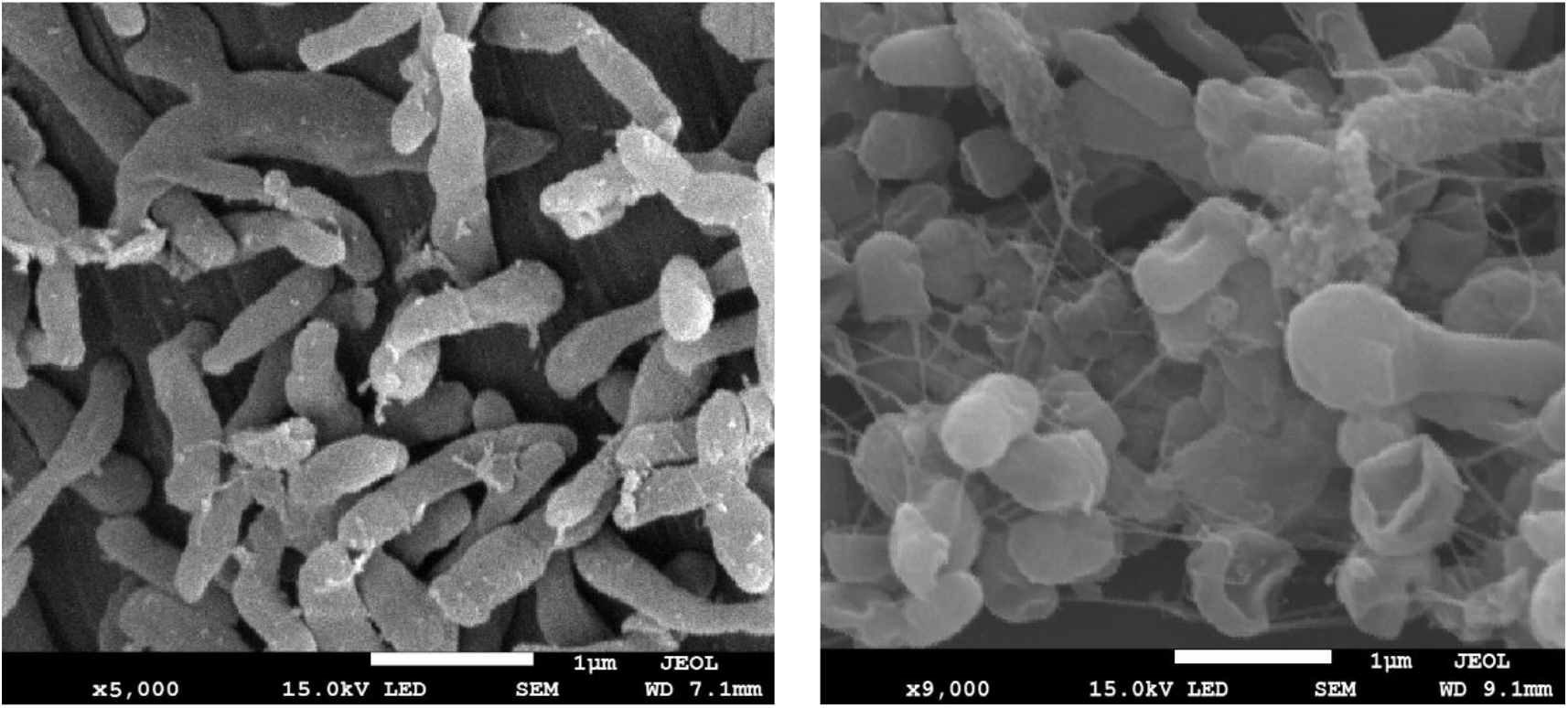
The microbial structure of *G.metallireducens* in anode biofilm

Fig.7 illustrates the electron transfer in the biofilm of the MFC electrode. The *G.metallireducens* of biofilm did not generate nanowires in MFC with electronic mediators, microbes in the inner layer of biofilm transmit electrons to the electrodes through direct contact, microbes in the outer layer of biofilm or solution transport electrons to the surface of electrodes through electronic mediators. The *G.metallireducens* of biofilm would generate nanowires in MFC without electronic mediators, microbes in the inner and outer layers of the biofilm are connected by nanowires to transfer electrons to the electrodes efficiently (Telma et al, 2015).

**Fig.7.**
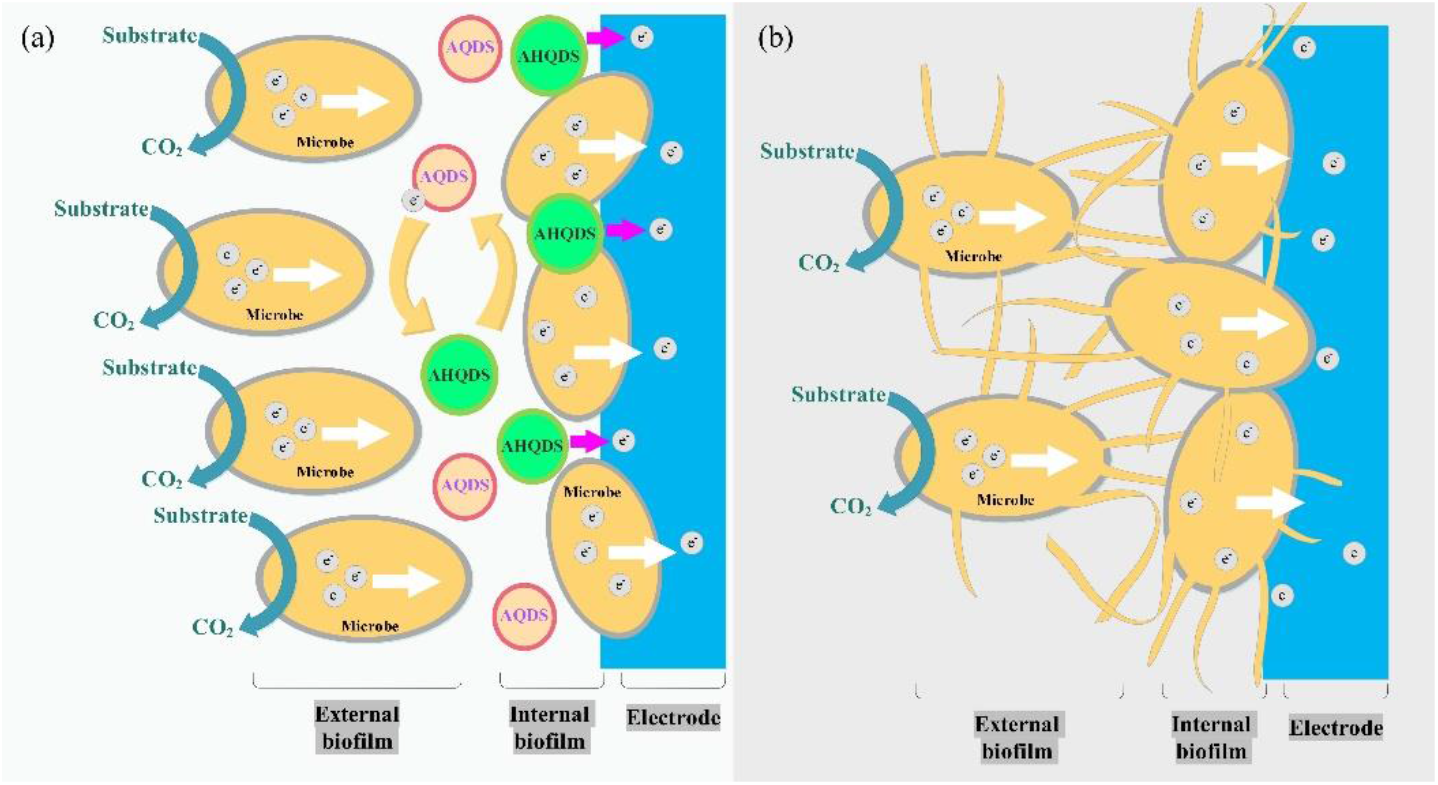
Electron transfers pathways for *G.metallireducens* in MFC.

### 3.3 Electrochemical characteristics of nanowire biofilm in MFC

Cyclic voltammetry (CV), electrochemical impedance spectrum (EIS), and linear step chronoamperometry (LSC) were used to characterize the electrochemical activity of the biofilm electrode. Fig.8a showed the CV profiles of ordinary biofilm electrodes (with AQDS) and nanowire biofilm electrodes (without AQDS). In a mixed buffer solution, a pair of redox peaks were observed clearly in the respective electrodes. But the nanowire biofilm electrodes showed higher CV peak current and smaller cathodic/anodic peak separation, indicating nanowire could increase the electrochemical activity of electrodes. The nanowire can accelerate the electron transfer between the microbes with the electrodes. The EIS plot consists of a semicircle (high-frequency region) and a straight line (low-frequency region). The diameter of the semicircle represents charge-transfer resistance (R_ct_), which value is negatively correlated with the electron transfer speed. As shown in Fig.8b, the semicircle of nanowire biofilm electrodes is smaller than that of ordinary biofilm electrodes, suggesting lower interfacial charge-transfer resistance (Rct) and a greater charge transfer rate. Linear step chronoamperometry is used to detect the electron transfer activity of the electrode. The results are shown in Fig.8c. Nanowire biofilm electrodes showed a stronger and broader current response than ordinary biofilm electrodes, which indicates nanowire biofilm electrodes could conduct stronger current under the same voltage condition. Here, it seemed to limit the metabolism of the biofilms has a similar impact as added to AQDS. This could be a consequence of the dominating electron mediator of the AQDS, changed diffusion regimes, or increased charge transfer resistance due to reduced metabolic activity. The above results suggested that the nanowire could significantly enhance the charge transfer rate and electrode current signal. The output voltage and performance of the MFC were improved.

**Fig. 8.**
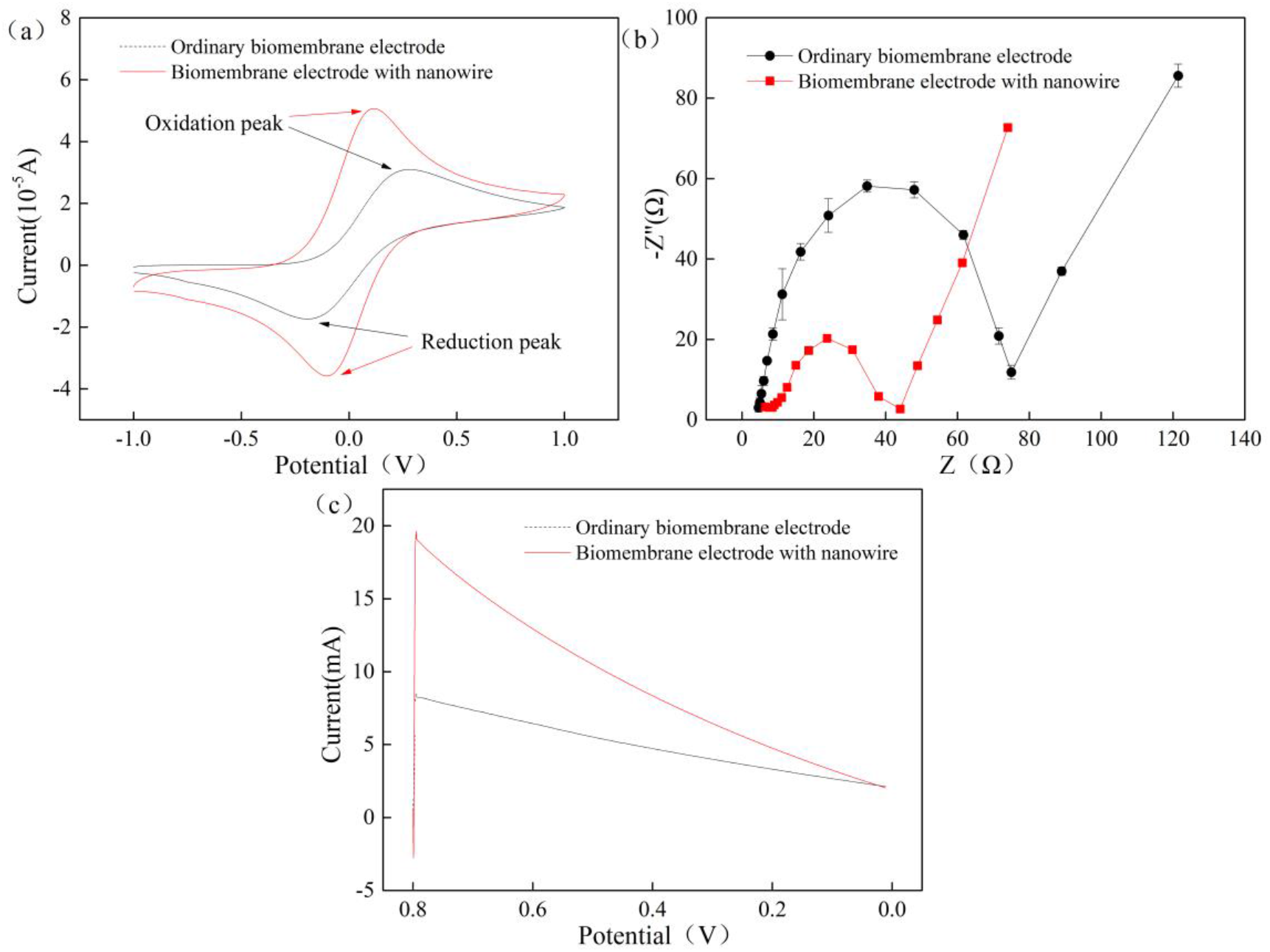
(a) CV of the ordinary biofilm electrode (dominated biofilms at constant electron mediator concentration (50μmol L–1 AQDS)) and nanowire biofilm electrode (dominated biofilms electrodes at the without electron mediator (AQDS)) in a mixed buffer (10 mmol/L phosphate+10 mmol/L NaAC) at scan rate of 5 mV/s, respectively; (b) EIS of the ordinary biofilm electrode and nanowire biofilm electrode in K_3_Fe(CN)_6_ (5 mmol/L)+K_4_Fe(CN)_6_ (5 mmol/L)+KCl (0.1 mol/L) solution (Frequency range of 0.1 Hz to 100 kHz with an AC signal of 5 mV amplitude); (c) LSC plots of the ordinary biofilm electrode and nanowire biofilm electrode (0.8~0 V and with a scan rate of 0.01 mV/s).

## 3 Conclusion

Compared with using Fe (OH)_3_ as a microbial electron acceptor directly, the addition of AQDS can accelerate the dissimilatory reduction of Fe (OH)_3_ in a preliminary stage. But the addition of AQDS hinders the growth of microbial nanowires, limits the upper bound of electron transfer efficiency. The accelerating effect of AQDS on microbial reduction was not significant after a long reaction time. By comparing microbial structure on electrode biofilm, the *G.metallireducens* did not generate nanowires with AQDS, microbes adhere to the graphite fibers of electrodes such as granules. And the *G.metallireducens* generated nanowires without AQDS. Nanowires are interlaced to connect microbes in biofilm, or microbes adhere to graphite fibers through nanowires. Microbial nanowires could significantly enhance the charge transfer rate and improve the performance of the MFC.

## Acknowledgments

This research was supported by the China Ocean Mineral Resource R&D Association under Grant JS-KTHT-2019-01 and No.DY135-B2-15, Major science and technology program for water pollution control and treatment under Grant No.2015ZX07205-003, the National Natural Science Foundation of China under Grant No.21176242 and No.21176026.

